# Development of muti-agent-simulation models for intercellular communication via cytokines and extracellular matrices

**DOI:** 10.1101/2022.09.21.508950

**Authors:** Ken-ichi Inoue, Satoko Kishimoto, Tomoki Mogami, Shigeru Toyoda, Masanori Hariyama

## Abstract

**Background:** Intercellular communication is a critical innovation during multicellular organismal evolution. Cells release / receive cytokines and utilize them as intercellular signal entities. How a well-orchestrated communication emerges from individual cell behavior remains largely unknown. Here we abstracted the biological phenomenon and developed multi-agent-simulation to investigate the intracellular communication.

**Methods:** Two dimensional MAS platform was developed using Artisoc 4.2.1 standard software. We focused on intercellular communication via cytokines and extracellular matrices. Three agents, “cells”, “cytokines” and “extracellular matrices” were defined and the interaction rules among the agents were designed. Two different mathematical models of cytokine-gradient determination were tested: spatial derivative and temporal derivative models. As a case study, neutrophil swarming was modeled and the cell swarming was defined as an evaluation criterion. Moreover, a surgically injured mouse model and a fluorescent time-lapse imaging were used to observe neutrophil swarming.

**Results:** We performed multiple simulations with six different virtual conditions, changing multiple parameters simultaneously and randomly. After 400 simulations for each condition, we counted the number of successful trials (i.e. neutrophil swarming within 10000 steps). Spatial derivative model showed more successes compared to temporal derivative model. Among eight parameters randomly assigned, cell’s exploration speed by random walk most remarkably influenced on success rate of neutrophil swarming. In *in vivo* model, bone marrow derived neutrophils gather towards the clumps with various migration speed. The mode of approaching resembles to spatial derivative model, rather than temporal derivative model.

**Conclusions:** MAS could be a useful approach to investigate the emergence of intercellular communication during multicellular evolution. Neutrophil could adopt the spatial derivative model as a sensing mechanism of cytokine gradient.

## Introduction

Multicellular organisms developed intercellular communication system during evolution to achieve the sufficient body size. Unicellular organism such as Escherichia coli, had already acquired the ability to “swim” toward / away from attractants / repellents, respectively [1]. Dictyostelium, a slime mold or social amoebae, has been used as a model organism because of peculiar multicellular assembly [2]. Volvox carteri, a green algae forms spherical colonies that are composed of two differentiated cell types, flagellate somatic cells and germ cells. Volvox is a primitive multicellular organism which acquired multifunctional extracellular matrices (ECM) out of the cell wall of unicellular ancestor [3]. Multicellular organisms are combining these primitive functional components for intercellular communication but the significance of each component, especially in the context of mutual interactions among components, remains largely unknown.

Multi agent based simulation (MAS) has been utilized to investigate the complex behavior such as inflammation or tumor microenvironments [4]. In MAS, multiple agents autonomously move according to the simple assigned rules, and the defined interactions among agents or environments display emergent behaviors. Even though the system as a whole does not have a control center, it achieves some degree of orchestration.

In this study, we developed a MAS which mimics intercellular communication via cytokines and ECM and reproduced neutrophil swarming (acute inflammation) as an emergent behavior. Two fundamental questions were addressed through the MAS: How cells sense the directionality of cytokine signals? What is a critical parameter to achieve the neutrophil swarming?

## Methods

### Modeling of intercellular communication via MAS

We developed a novel two dimensional (2D) MAS platform using Artisoc 4.2.1 standard software (Kozo Keikaku Engineering Inc., Tokyo, Japan). In this prototype model, we adopted a simplified and abstracted description of biological phenomena, omitting intracellular signal cascades, nutrient metabolism, cell division and apoptosis. Instead, we focused on intercellular communication via cytokines and ECM. Cells, cytokines and ECM influence each other through biochemical reactions and change their status every moment (mutual interactions). Therefore we defined three agents, “cells”, “cytokines” and “ECM” and designed the interaction rules (algorithms) as follows.

### Definition of ECM agents

ECM are composed by collagens or laminins, produced or degraded by nearby cells. In MAS, 2D plane was divided into L x M grids (a grid represents 1 µm x 1 µm square dot) and each ECM density was defined as 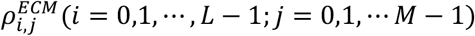.

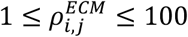

Here 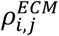 is a relative unit and the more 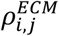, the more rigid the ECM in the particular grid is. Cytokines and cells move into and overlap with ECM grids. ECM influence on the behavior of cells because cells prefer more rigid scaffolds during migration. ECM also influence on the diffusion speed and protein stability of cytokines because freely available cytokines quickly diffuse and are under proteinase degradation. By binding to ECM, cytokines escape from the diffusion and degradation, increasing the stability and half-life.

### Definition of cytokine agents

Cytokines are intercellular signal entities released (produced) by cells and received (signal-sensed) by cells. In the MAS, particular cytokines that can influence on cellular motility (i.e. cells are attracted and migrate toward cytokines) were modeled and investigated. Two modes of cytokine behavior were defined, diffusion and degradation. Since cytokines are small molecules and follow Brownian motion, the diffusion of each cytokine was modeled as random walk of among the 2D grid. The direction of each step was defined as *θ* (uniformly distributed random number) and the distance of the random walk was inversely related to the square of ECM density. Here we approximated the ECM viscosity *η* to ECM density 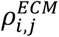 and the functional relationship between *d* and 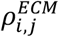 was determined based on Stokes-Einstein equation. Therefore, the random walk of each cytokine was defined as follows:

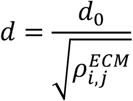

Here, *d*_0_(> 0) is a parameter to determine the distance of one step walk. *i,j* are coordinates among ECM grid in which the cytokine exists before the one step walk. Cytokine degradation (half-life) was modeled as the probability for each cytokine to disappear. The stability of cytokines increase when the density of 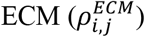 is higher (protection effect). Thus, the probability of cytokine disappearance was defined as follows:

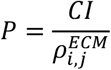

Here *CI* (0 ≤ *CI* ≤ 1) is a probabilistic parameter which determines “cytokine instability” (faster degradation). Cytokines do not collide with cells and occupy the same space simultaneously.

### Definition of cell agents

In this study, we selected neutrophils as a representative of stromal cells. In MAS, cell morphology is defined as 2D circle with the radius 15 µm. As a summary of agent behaviors (rules), we presented the flow-chart in Figure 1.

**Figure 1.**
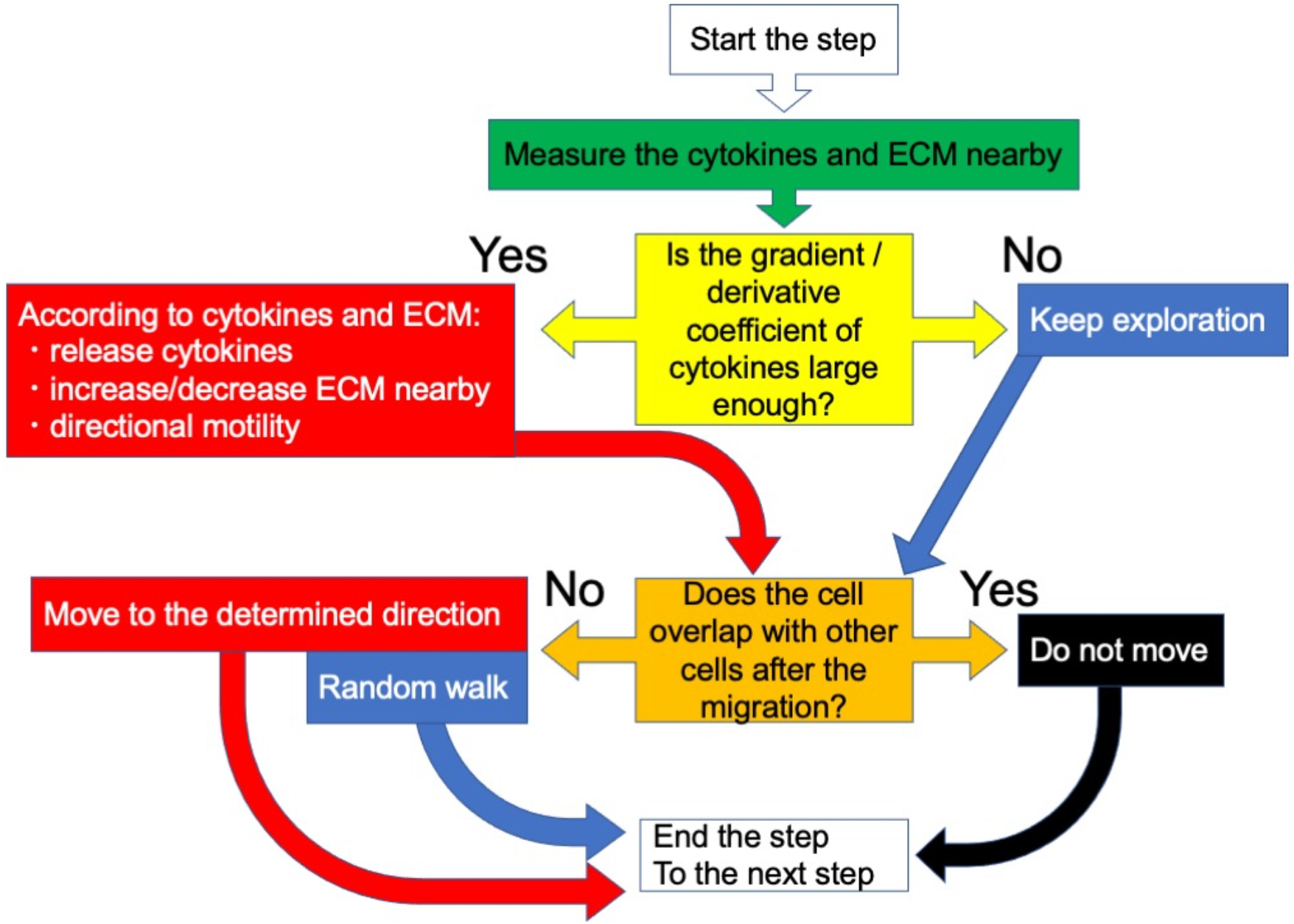
Flowchart of cell agents’ behaviors in each step. At the beginning of the step, each cell measure the concentration of cytokines and the density of ECM nearby (green color). If the gradient (in spatial derivative model) or derivative coefficient of (in temporal derivative model) of cytokines is larger than the threshold (yellow color), cells execute three functions: releasing cytokines, increasing/decreasing ECM, and move with certain directionality (red color). Otherwise, cells keep environmental exploration (blue color, random walk). Since cells occupy certain amount of space, they cannot overlap each other. Thus, if the cells overlap after directional motility or random walk, they cancel the movement and stay there (black color).

### Two distinct modes of cytokine-gradient determination: spatial derivative and temporal derivative models

We investigated two different hypotheses about underlying mechanism of cytokine-gradient determination, 1) Are cells attracted toward cytokine gradient by sensing / comparing cytokine concentration in all directions? 2) Do cells explore the environment by intensive random walk and sense the temporal difference of cytokines during the random walk? The former and latter hypotheses were investigated by two different mathematical models, spatial derivative and temporal derivative models, respectively.

#### 1. Spatial derivative model

A cell is represented by a circle with the radius *R* (Figure 2A). A cell measures the concentration of cytokines in both the center of the cell (Figure 2A, a concentric circle with the radius 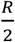) as well as around the surface of the cell (Figure 2A, circumferential circles in eight-directions with the radius 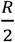). The cytokine concentration inside the particular circle is defined as:

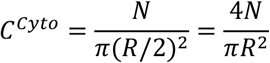

**Figure 2.**
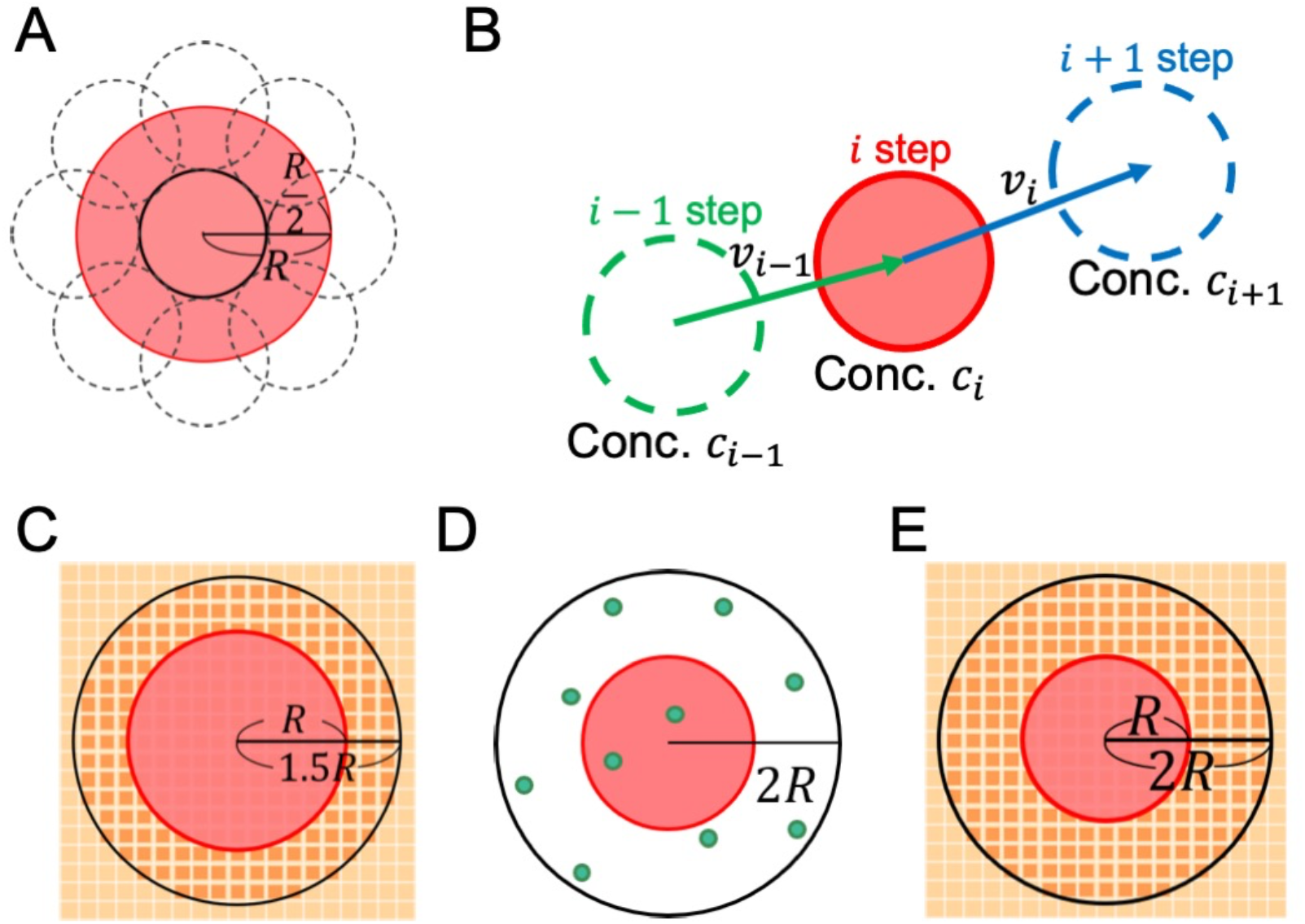
Schematic representation of two modes of cytokine gradient determination and interaction rules among cells, ECM and cytokines. A. Spatial derivative model A cell is represented in red-highlighted circle with the radius *R*. A cell measure the concentration of cytokines in both the center of the cell (presented as a solid line circle, with the radius 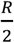) as well as around the surface of the cell (eight broken line circles with the radius 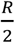, in eight-directions), followed by approximate determination of the concentration gradient. B. Temporal derivative model In temporal derivative model, cytokine gradient is determined based on the difference between cytokine concentration of *i* step and *i* − 1 step. The concentration is calculated, counting cytokine number overlapping with the cell (i.e., the circle with the radius *R*). 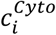 : concentration of cytokines overlapping with the cell (within the circle) at *i* step. *v*_*i*−1_: the distance of cell migration before *i* step. C. A cell (red-highlighted circles) measures all the ECM densities (dark orange) within a concentric circle (black), from that a mean ECM density is calculated as an average. D. A cell releases cytokines (right green circles) randomly within a concentric circle with the radius 2*R*. E. A cell changes local ECM density in every grids within a concentric circle with the radius 2*R* (dark orange squares).

Here *N* represents the number of cytokines within the area.

The cytokine gradient in *i* step is defined as follows: First the *C*^*Cyto*^ is measured inside the cell center (solid line circle in Figure 2A) and termed as 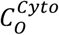. Next the *C*^*Cyto*^ is measured around the cell surface (eight broken line circles in Figure 2A) and the maximum concentration is termed as 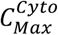. Subsequently the cytokine gradient is calculated as:

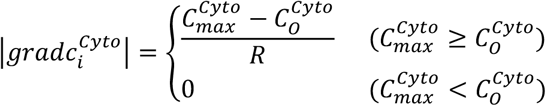

The direction of 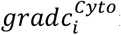 is determined from the center of the cell to the direction in which the 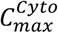 is given. If more than two identical 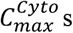 are given, one direction is determined using uniformly distributed random number. A mean cytokine concentration around the cell 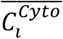 is calculated by averaging *C*^*Cyto*^ in eight directions at *i* step. This model is based on the biological assumption that cells have a kind of cytokine sensor and measure the concentration in eight different directions.

#### B. Temporal derivative model

In temporal derivative model, cytokine gradient is determined based on the difference between cytokine concentration of *i* step and *i* − 1 step (Figure 2B). The concentration is calculated, counting cytokine number overlapping with the cell (i.e., the circle with the radius *R*).

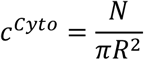

Suppose the cytokine concentration at *i* step as 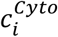 and the distance of cell migration as *v*_*i*−1_ (Figure 2B), then the cytokine gradient is defined as follows:

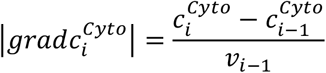

The direction of 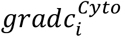 is determined based on *θ*_*i*−1_ (direction of cell’s random walk at one step earlier) as follows:

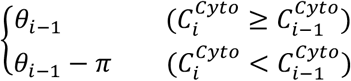

Namely, cells sense the increase 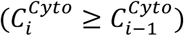 or decrease 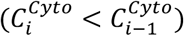 of cytokines during random walk and roughly estimate whether they are going to right or wrong direction for haptotaxis. Cells turn to opposite (*θ*_*i*−1_ − *π*) if they are going wrong direction, seeking the more cytokines.

### Measurement of ECM by cells

Mean density of 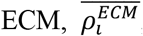, is calculated as an average of 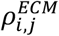 within a surrounding circle with a radius of 1.5*R* (Figure 2C).

#### Haptotaxis

Haptotaxis is the directional motility upward a gradient of cytokines bound to 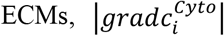. Since cells utilize ECM as solid scaffold during migration, higher 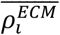 is preferred.

Therefore, the moving distance is defined:

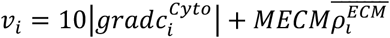

Here *MECM* (mobility in ECM, ≥ 0) indicates a parameter how much the mobility is affected by ECM.

### Cytokine production by cells

Releasing cytokines by cells is enhanced when the cytokine gradient exceeds the threshold (signal augmentation), but it is suppressed when the mean cytokine concentration gets high (negative feedback). A cell releases cytokines randomly within a concentric circle with the radius 2*R* (Figure 2D). A number of cytokine production is defined as follows.

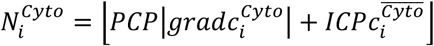

Here *PCP* (promotion of cytokine production, ≥ 0) or *ICP* (inhibition of cytokine production, < 0) indicates a parameter which positively or negatively influences on cytokine production, respectively. ⌊*x*⌋ denotes dropping the fractional portion of the number. When 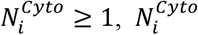 cytokines are released.

### Production / degradation of ECM by cells

Cells increase ECM density nearby when the cytokine gradient exceeds the threshold (response to signal), but it decrease ECM density when mean local ECM density gets high (negative feedback). After measuring 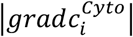 and 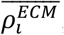, a cell changes local 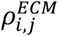 in every grids within a concentric circle with the radius 2*R* (Figure 2E). The amount of change in each grid is defined as follows:

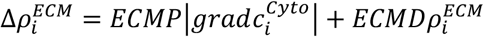

Here *ECMP* (ECM production, ≥ 0) or *ECMD* (ECM degradation, < 0) indicates a parameter which positively or negatively influences on ECM change, respectively. When 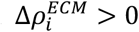, ECM are produced, but when 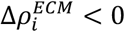, ECM are degraded.

### Environmental exploration of cells by random walk

When a cytokine gradient is not high enough, cells cease haptotaxis and switch to exploratory random walk. Similar to the random walk of cytokines, the direction of each step was defined as *θ* (uniformly distributed random number). And the distance of migration *v*_*i*_ is defined:

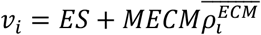

Here *ES* (exploratory speed, ≥ 0) indicates a parameter.

### Evaluation of MAS

We performed a middle scale 2D MAS (450 µm x 450 µm or 450 × 450 ECMs grids) to recapitulate neutrophil swarming *in silico* (Figure 3A). At the start of MAS, eight cells evenly distributed within the 2D space. Inflammation epicenter is placed, releasing ten cytokines per step, eventually 25000 cytokines throughout 5000 steps.

**Figure 3.**
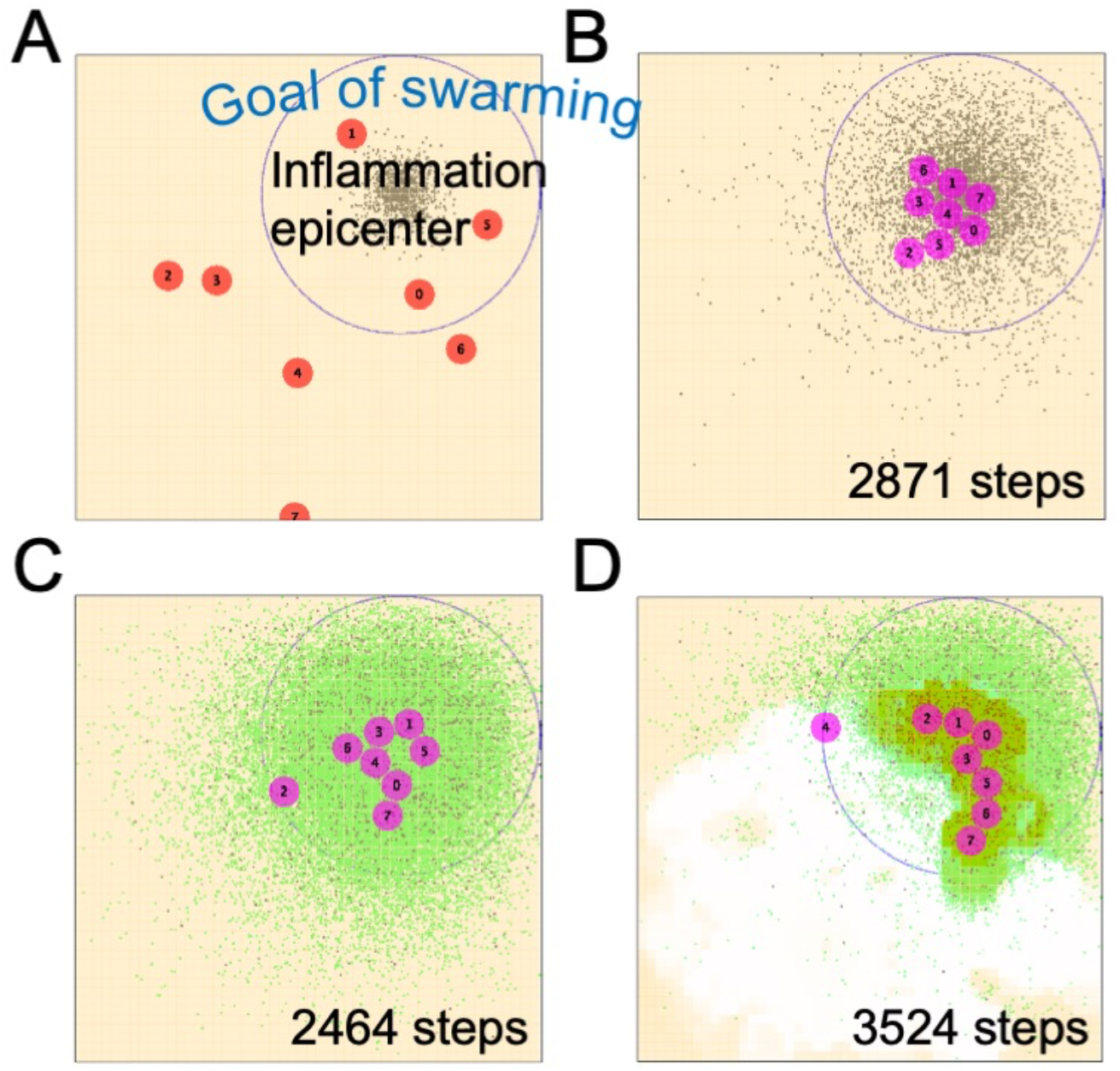
2D MAS of neutrophil swarming with different experimental conditions. A. MAS with the size of 450 µm x 450 µm (450 × 450 ECMs grids). Eight cells (red circles) are numbered and placed randomly. Cytokines (black dots) are continuously released from “inflammation epicenter”, mimicking tissue damage or pathogen infection. Goal of swarming is set surrounding the epicenter (blue empty circle). Neutrophil swarming is complete once all the eight cells go across the goal (B, C and D). B. Neutrophil swarming with condition 1 (without cytokine release, without ECM production / degradation, spatial derivative). Cells change their color from red to pink, when they receive cytokine signal (i.e. the cytokine gradient exceeds the threshold). Accordingly cells change the mode of migration from random walk to directional motility. C. Neutrophil swarming with condition 2 (with cytokine release, without ECM production / degradation, spatial derivative). Green dots indicate newly produced cytokines released from cells. D. Neutrophil swarming with condition 3 (with cytokine release, with ECM production / degradation, spatial derivative). Note that ECM (pale orange background) is produced or degraded due to ECM remodeling by cells.

Neutrophil swarming is defined as all the eight cells gather around the inflammation epicenter (within a defined circle, Figure 3A). An experiment was counted as successful when neutrophil swarming happened within the first 10000 steps. An experiment was terminated after 10000 steps if the neutrophil swarming did not happen and was counted as a failure.

### Animals and *in vivo* imaging

C57BL/6-Tg(Acta2-DsRed)1Rkl/J mice, which express DsRed red fluorescent protein (RFP) under the alpha smooth muscle actin (αSMA) promoter, from the Jackson Laboratory (Bar Harbor, ME) and C57BL/6-Tg(CAG-EGFP) mice, which express enhanced green fluorescent protein (GFP) from Japan SLC, Inc. (Hamamatsu, Japan) bred and maintained by the Research Center for Laboratory Animals, Dokkyo Medical University. All animal experiments adhered to the Guidelines for Animal Experimentation of Dokkyo Medical University, with all efforts made to minimize animal numbers and suffering. To establish a mouse model that displays GFP immune cells and RFP-activated myofibroblasts, bone marrow transplantation (BMT) was performed. Recipient 10- to 12-wk-old Acta2-DsRed mice were whole body-irradiated at 9 Gy per mouse (MBR-1520R-4, Hitachi Medical Co. Ltd., Tokyo, Japan). Donor GFP mice were euthanized, and bone marrow cells from femur were flushed with Dulbecco’s Modified Eagle Medium (DMEM; Life Technologies Oriental, Tokyo, Japan) containing 2% (vol/vol) Fetal Bovine Serum (FBS). The suspension was passed through a 100-µm strainer (Greiner Bio-One, Frickenhausen, Germany), centrifuged at 490 g for 5 min (EX-136; Tomy Seiko Co., Ltd., Tokyo, Japan), and resuspended in saline. Within one hour after the radiation, recipient mice received 5-10 × 10^6^ donor cells via tail-vein injection. Following BMT, mice received enrofloxacin (Baytril^®^10% solution, Bayer, Germany) in the water for 2 weeks. Control mice that were radiated but not transplanted died within 12-13 d.

Time lapse *in vivo* imaging of mouse was taken one month after the BMT. Mouse was anesthetized with 4% isoflurane and maintained at 1.5-2%. On the day before the observation, the epididymal fat of the mouse was surgically injured by mincing the fat parenchyma and cauterizing the fat-feeding artery (Stem Cells Int., Kishimoto, et al. 2020). Injured epididymal fat was exposed outside of the mouse body and occasionally moistened with saline. Time-lapse images were captured at 30-second intervals using 20× dry (NA = 0.45) objectives mounted on a motorized Keyence BZ-X810 (Osaka, Japan) fluorescence microscope and image data exported using BZ-X800 Analyzer to create movies.

## Results

### A case study of MAS: neutrophil swarming as an emergent behavior

Here We performed multiple simulations with six different virtual conditions, changing multiple parameters (*d*_0_, *CI, MECM, PCP, ICP, ECMP, ECMD, ES*) simultaneously and randomly.

Condition 1: without cytokine release, without ECM production / degradation, spatial derivative (Figure 3B)

Condition 1’: without cytokine release, without ECM production / degradation, temporal derivative

Condition 2: with cytokine release, without ECM production / degradation, spatial derivative (Figure 3C)

Condition 2’: with cytokine release, without ECM production / degradation, temporal derivative

Condition 3: with cytokine release, with ECM production / degradation, spatial derivative (Figure 3D)

Condition 3’: with cytokine release, with ECM production / degradation, temporal derivative

After 400 simulations for each condition, we counted the number of successful trials (i.e. neutrophil swarming within 10000 steps). Overall, spatial derivative model showed more successes compared to temporal derivative model (117 vs. 65 for condition 1 vs. 1’, 166 vs. 96 for condition 2 vs. 2’, 115 vs. 59 for condition 3 vs. 3’, *P* = 0.008 by paired-*t* test, Table1). In spatial derivative model, cells aggregate each other (Figure 3B, C and D) and did not move after that. Releasing cytokines by cells augmented the signal and accelerated the swarming (117 vs. 166 for condition 1 vs. 2, Figure 3B and C). On the other hand, ECMs production/degradation delayed the swarming (166 vs. 115 for condition 2 vs. 3, Figure 3C and D).

**Table 1.** Success ratio of neutrophil swarming.

| Spatial differentiation |  |  |  | Time derivative |  |  |  |
| --- | --- | --- | --- | --- | --- | --- | --- |
|  | Total trials | Successes | Ratio |  | Total trials | Successes | Ratio |
| A | 400 | 117 | 29.25% | A' | 400 | 65 | 16.25% |
| B | 400 | 166 | 41.50% | B' | 400 | 96 | 24.00% |
| C | 400 | 115 | 28.75% | C' | 400 | 59 | 14.75% |

MAS were performed 400 times for each experimental condition.

Among eight parameters randomly assigned, cells’ exploration speed by random walk (*ES*) most remarkably influenced on success rate of neutrophil swarming (Figure 4). In both spatial derivative and temporal derivative models, *ES* proportionally increased success rate of the swarming. Data indicate that the speed of cells’ random expedition is a most critical parameter for efficient emergence of neutrophil swarming.

**Figure 4.**
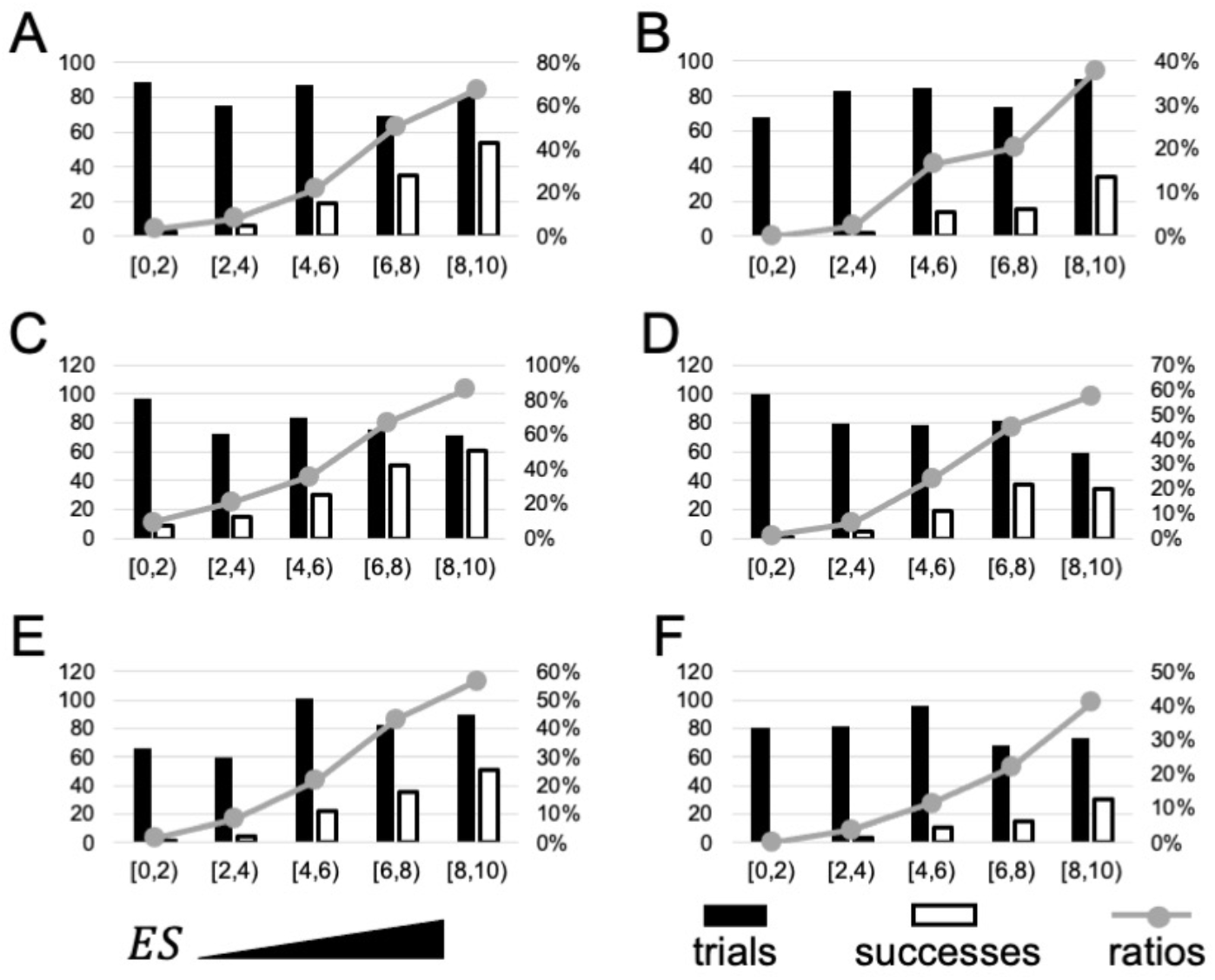
Exploratory speed of cells proportionately increase the success rate of neutrophil swarming. Histograms of MAS trials according to a parameter *ES*. Black and white bars indicate total trials and number of successes, respectively. Gray line graphs indicate the ratio of success. A. Condition 1: without cytokine release, without ECM production / degradation, spatial derivative. B. Condition 1’: without cytokine release, without ECM production / degradation, temporal derivative. C. Condition 2: with cytokine release, without ECM production / degradation, spatial derivative. D. Condition 2’: with cytokine release, without ECM production / degradation, temporal derivative. E. Condition 3: with cytokine release, with ECM production / degradation, spatial derivative. F. Condition 3’: with cytokine release, with ECM production / degradation, temporal derivative. *ES*: exploratory speed.

### Time lapse *in vivo* imaging of bone marrow derived cells after tissue injury

We observed bone marrow derived neutrophil mobilization in surgically injured mouse epididymal fat. One day after tissue injury, clumps of GFP positive cells were observed (Figure 5, Supplementary Movie 1). Interestingly, several GFP positive cells gather towards the clumps with various migration speed (Figure 5, Supplementary Movie 1). The mode of approaching resembles to spatial derivative model, rather than temporal derivative model, suggesting the hypothesis that neutrophil adopt the former as a sensing mechanism of cytokine gradient.

**Figure 5.**
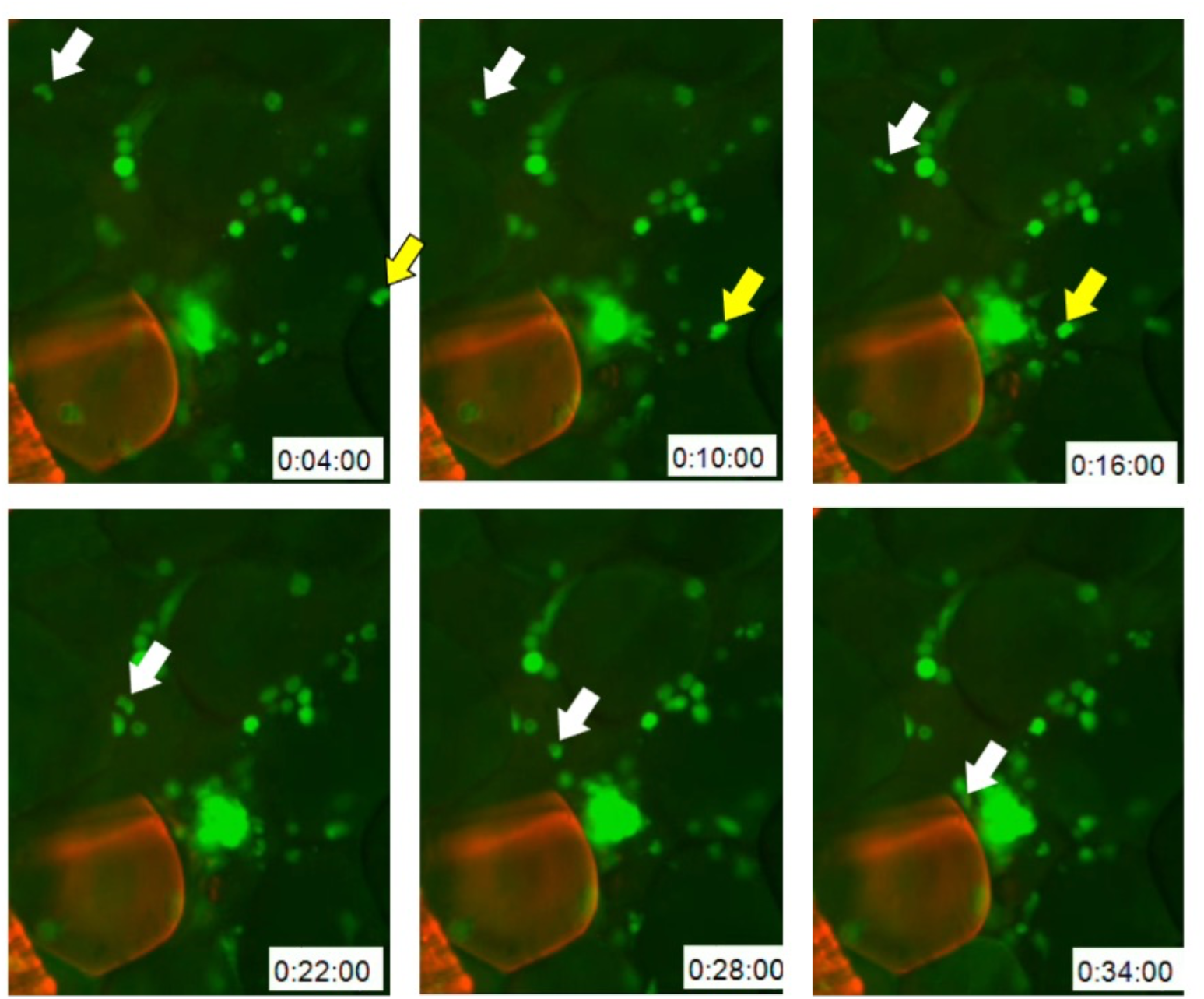
Time lapse *in vivo* imaging of bone marrow derived neutrophils swarming into cell clump. Fluorescent microscopic images were serially taken every four minutes in the anesthetized mouse epididymal fat. The fat was surgically injured one day before the observation, GFP positive cells were recruited from the bone marrow. Note that the cell with white arrow or the yellow arrow approaches to the cell clumps with different speed (the yellow is faster than the white). Red fluorescence label smooth muscle cells surrounding arteriole by *Acta2* promoter-driven DsRed.

## Discussion

Stromal cells such as tissue resident fibroblasts and macrophages cooperatively participate in tissue homeostasis [5, 6]. During pathogen infection or tissue damage, the tissue resident cells sense the emergency and transmit the signal via intercellular signal ligands such as cytokines [7]. As a results, the resident cells recruit immune cells from bone marrow, displaying chronic inflammation locally and transiently [8, 9]. The immediate inflammatory response is necessary for efficient tissue repair such as angiogenesis and parenchymal regeneration [10].

In this study, we abstracted the phenomenon, intercellular communication via cytokines, and developed MAS. Changing / applying multiple parameters in the MAS is corresponding to trial and error (optimization) in Darwinian evolution. By setting particular performance as a fitness variable, we can demonstrate “the selection of the fittest” theory *in silico*. As a case study, we picked up neutrophil swarming as a fitness variable. Two findings we obtained.

We investigated two different mechanisms of cytokine-gradient determination, spatial derivative and temporal derivative models, respectively. After extensive parameter search, we concluded that the spatial derivative is more suitable for neutrophil swarming (Table 1). Importantly, behaviors of neutrophils in the injured mouse tissue were similar to the spatial derivative model (Figure 5, Supplementary Movie1), suggesting that the mammalian neutrophils are adopting the “spatial sensors” to determine the movements.

Another important finding is that the speed of random exploration (cell movement) is critical in both spatial and temporal derivative models (Figure 4). Intercellular communication of Social amoebae utilize cAMP instead of cytokines [2]. Both cAMP and cytokines are not suitable materials for remote communication because the diffusion speed is too slow. Indeed, the passive diffusion *per se* were not sufficient to cover all the investigated area and active exploration of cells are critical to achieve the swarming (Figure 3B).

Releasing cytokines from each cell was not necessary but advantageous for swarming (Figure 3B and C, Table 1). Cytokine release enables “cytokine relay” among cells, expanding the area of cytokine signal from the epicenter (Figure 3C). Thus, autonomous cell exploration cells plus cytokine relay accelerate the swarming and increased the fitness. However, this could be disadvantageous from another aspect. When we consider the aspect of inflammation resolution, excessive augmentation of cytokine signal could lead to “brake failure”. To address such trade-off question, future research should include negative feedback mechanism for the cytokine release.

A limitation of this study is that the significance of ECM remains obscure from the MAS. ECM influences not only on cell movement, but also cytokine stability and diffusion speed. In the case study of neutrophil swarming, ECM synthesis / degradation rather decreased the fitness (Figure 3C and D, Table 1). However, this could not be the case if we model the fitness in different perspective, such as fibroblast behavior. To address the question about ECM significance, we need extension of cell agents’ behavior such as proliferation and apoptosis, mimicking the sequential tissue repair processes [10]. Equally important extension is the number of different cell types in the MAS. In tissue homeostasis, several different cell types constitute the cellular society [5, 6], necessitating including different set of rules for cell agents. In the future, we expand the MAS and address the different aspect of tissue repair such as efferocytosis and inflammation resolution.

## Acknowledgements

This study is funded by JSPS KAKENHI [grant number 21K12651].

## References

1. Wadhams GH, Armitage JP. Making sense of it all: Bacterial chemotaxis. Nature Reviews Molecular Cell Biology. 2004;5(12):1024–37. doi: 10.1038/nrm1524. PubMed PMID: WOS:000225711100015.

2. Artemenko Y, Lampert TJ, Devreotes PN. Moving towards a paradigm: common mechanisms of chemotactic signaling in Dictyostelium and mammalian leukocytes. Cellular and Molecular Life Sciences. 2014;71(19):3711–47. doi: 10.1007/s00018-014-1638-8. PubMed PMID: WOS:000341909100005.

3. Hallmann A. Extracellular matrix and sex-inducing pheromone in Volvox. International Review of Cytology - a Survey of Cell Biology, Vol 27. 2003;227:131-+. doi: 10.1016/s0074-7696(03)01009-x. PubMed PMID: WOS:000185673500003.

4. Yu JS, Bagheri N. Agent-Based Models Predict Emergent Behavior of Heterogeneous Cell Populations in Dynamic Microenvironments. Frontiers in Bioengineering and Biotechnology. 2020;8. doi: 10.3389/fbioe.2020.00249. PubMed PMID: WOS:000543893300001.

5. Plikus MV, Wang XJ, Sinha S, Forte E, Thompson SM, Herzog EL, et al. Fibroblasts: Origins, definitions, and functions in health and disease. Cell. 2021;184(15):3852–72. doi: 10.1016/j.cell.2021.06.024. PubMed PMID: WOS:000676120800005.

6. Haldar M, Murphy KM. Origin, development, and homeostasis of tissue-resident macrophages. Immunological Reviews. 2014;262(1):25–35. doi: 10.1111/imr.12215. PubMed PMID: WOS:000343997600003.

7. Bagnall J, Boddington C, England H, Brignall R, Downton P, Alsoufi Z, et al. Quantitative analysis of competitive cytokine signaling predicts tissue thresholds for the propagation of macrophage activation. Science Signaling. 2018;11(540). doi: 10.1126/scisignal.aaf3998. PubMed PMID: WOS:000439579100001.

8. Kishimoto S, Inoue K, Nakamura S, Hattori H, Ishihara M, Sakuma M, et al. Low-molecular weight heparin protamine complex augmented the potential of adipose-derived stromal cells to ameliorate limb ischemia. Atherosclerosis. 2016;249:132–9. doi: 10.1016/j.atherosclerosis.2016.04.003. PubMed PMID: WOS:000376505800020.

9. Kishimoto S, Inoue K, Sohma R, Toyoda S, Sakuma M, Inoue T, et al. Surgical Injury and Ischemia Prime the Adipose Stromal Vascular Fraction and Increase Angiogenic Capacity in a Mouse Limb Ischemia Model. Stem Cells International. 2020;2020. doi: 10.1155/2020/7219149. PubMed PMID: WOS:000538840100001.

10. Eming SA, Wynn TA, Martin P. Inflammation and metabolism in tissue repair and regeneration. Science. 2017;356(6342):1026–30. doi: 10.1126/science.aam7928. PubMed PMID: WOS:000402871700041.

